# Perceptual Gravity Weighting Is Associated with Cybersickness Susceptibility During Virtual Reality

**DOI:** 10.64898/2026.07.31.741852

**Authors:** Megan H. Goar, Michael Barnett-Cowan

## Abstract

Cybersickness in virtual reality (VR) arises from conflicts between sensory signals, yet susceptibility varies considerably across individuals. Previous work in this cohort demonstrated that vestibulomotor responses during postural control are associated with cybersickness susceptibility. Here, we examined whether perceptual weighting of gravity, visual, and body cues used to estimate upright orientation is similarly associated with cybersickness and related to previously reported vestibulomotor measures. Thirty-eight healthy young adults (21 females, 17 males) completed a standing VR rollercoaster task while receiving continuous stochastic electrical vestibular stimulation (0–25 Hz, ±4.5 mA). In the current analysis, perceptual cue weights were quantified before and after VR using the Oriented Character Recognition Task. Cybersickness was assessed using the Fast Motion Sickness Scale (FMS), and participants were classified as non-sick (FMS < 5), medium-sick (FMS ≥ 5), or high-sick (terminated the VR exposure early due to intolerance). Before VR, non-sick participants exhibited greater gravity weighting (36% vs. 25%) than high-sick participants, whereas high-sick participants showed a non-significant trend toward greater visual weighting (30% vs. 18%). Perceptual cue weights changed minimally following VR, and the magnitude of perceptual reweighting was not associated with sickness severity. Vestibulomotor measures were not correlated with perceptual gravity weighting, and postural sway during VR was not associated with visual weighting. These findings suggest that greater baseline perceptual gravity weighting, rather than short-term perceptual reweighting, is associated with reduced cybersickness susceptibility. The dissociation between perceptual and vestibulomotor measures suggests that orientation perception and postural control reflect partially distinct multisensory integration processes.

**New and Noteworthy:** This study demonstrates that baseline perceptual gravity weighting is associated with susceptibility to cybersickness during virtual reality exposure with concurrent electrical vestibular stimulation. Greater reliance on gravity cues was associated with reduced susceptibility, whereas visual weighting showed a similar but non-significant trend and perceptual reweighting changed minimally following exposure. Perceptual measures were dissociable from vestibulomotor responses, suggesting that orientation perception and postural control reflect partially distinct multisensory integration processes.

## Introduction

The central nervous system (CNS) maintains upright orientation and postural stability by integrating vestibular, visual, and somatosensory inputs (1). Under typical conditions, these sensory signals are largely congruent and enable accurate perception of body orientation relative to gravity (1). When the reliability of these inputs changes, creating a mismatch between expected and incoming afferent information, the CNS can flexibly reweight their relative contributions to resolve conflict and thereby maintain stable perception and postural control.

Sensory reweighting is thought to occur during immersion in virtual reality (VR), where visual displays simulate self-motion in the absence of corresponding vestibular or proprioceptive input (2–4). Cybersickness, a visually induced form of motion sickness, is a common consequence of this sensory conflict and is characterized by symptoms such as nausea, headache, and vomiting (12–18). Despite its widespread applications in training, rehabilitation, education, and entertainment, the use of VR in fields such as aviation, surgery, rehabilitation, and education remains limited by cybersickness (6,7,9,10). Evidence suggests that the extent of sensory reweighting is inversely related to cybersickness susceptibility (11,12), which varies considerably across individuals (11,13).

Recent findings suggest a more complex relationship between sensory reweighting and cybersickness susceptibility. Using stochastic electrical vestibular stimulation (EVS) and medial-lateral center-of-pressure (ML-CoP) responses, we previously demonstrated that lower baseline vestibulomotor coupling was associated with reduced susceptibility to cybersickness, whereas dynamic downweighting of vestibular input occurred over time only in individuals who developed cybersickness (14). However, this adaptive reweighting was insufficient to prevent increases in postural sway and symptom progression, suggesting that baseline sensory weighting may play a more prominent role than reweighting capacity alone in determining cybersickness susceptibility. These findings, however, were limited to vestibulomotor pathways involved in postural control. Whether baseline perceptual sensory weighting and its adaptation during VR similarly relate to cybersickness susceptibility, or correspond with previously identified vestibulomotor markers, remains unknown.

Perceptual measures provide a complementary perspective on multisensory integration by probing how the CNS combines sensory cues to generate conscious judgments about orientation. One commonly used perceptual measure is the subjective visual vertical (SVV), a psychophysical method to assess perceptual alignment with gravity (15–18). In this task, participants judge whether a visual line aligns with their perceived direction of gravity across different body orientations and visual backgrounds, reflecting the relative contributions of gravity, visual, and body (idiotropic) cues to orientation perception. Chung & Barnett-Cowan (2023) found that individuals who exhibited greater shifts in SVV following VR exposure were less susceptible to cybersickness, suggesting successful perceptual reweighting. In this study, baseline perceptual weighting was not found to predict susceptibility.

A related paradigm, the Oriented ChAracter Recognition Task (OCHART), similarly estimates the relative contributions of gravity, body, and visual cues to perceived upright orientation (16–18). In this task, participants identify whether a briefly presented character is a “p” or a “d” across many trials. Unlike the SVV, which reflects the perceived direction of gravity, the OCHART assesses judgments about the orientation of external objects. Importantly, OCHART appears particularly sensitive to visual influences, making it well suited for investigating perceptual estimates of upright in visually driven environments such as VR.

To date, no study has examined whether perceptual estimates of upright correspond with previously identified vestibulomotor markers of cybersickness within the same individuals. Consequently, it remains unclear whether perceptual and sensorimotor indices of sensory weighting reflect shared mechanisms of adaptation to sensory conflict or partially distinct multisensory integration processes. Additionally, it is unknown whether individuals exhibiting greater visual weighting in perceptual judgments also demonstrate greater postural sway during VR exposure, as greater reliance on visual information may increase susceptibility to visually induced perturbations and, consequently, visually driven postural responses. The present study addressed these questions by examining perceptual sensory weighting before and after VR exposure using the OCHART and relating these measures to cybersickness severity and previously characterized vestibulomotor responses.

We hypothesized that greater reliance on gravity cues and reduced reliance on visual cues would be associated with reduced susceptibility to cybersickness because individuals who rely less on visual information may be less influenced by the visually induced sensory conflict encountered during VR. We further examined whether perceptual sensory weighting paralleled vestibulomotor responses and whether postural sway during VR exposure was associated with perceptual visual weighting. If shared mechanisms underlie perceptual and vestibulomotor sensory weighting, individuals exhibiting stronger vestibulomotor coupling would also be expected to exhibit greater gravity weighting and reduced visual weighting during perceptual orientation judgments. Conversely, a lack of correspondence would suggest that perceptual orientation and postural control rely on partially independent multisensory integration processes. Finally, we hypothesized that individuals with greater visual weights would exhibit larger postural sway responses during VR exposure, reflecting greater susceptibility to visually induced perturbations. Understanding how perceptual sensory weighting relates to cybersickness susceptibility may advance our understanding of multisensory integration and inform strategies to improve user experience in virtual environments.

## Methods

### Participants and Ethical Approval

A total of 38 participants (21 females, 17 males) were recruited as part of a larger study investigating vestibulomotor and perceptual sensory weighting during VR exposure. The present study represents a secondary analysis of this cohort focused on perceptual sensory weighting. The sample size was determined *a priori* for the original study based on the primary vestibulomotor outcomes. A power analysis for a two-tailed repeated-measures *t*-test, assuming a power of 0.80, α = 0.05, and an effect size of Cohen’s *d* = 1.0, indicated that 10 participants were required per group. With three sickness groups, the target sample size was therefore 30 participants. To account for participant withdrawal, equipment malfunctions, and other unforeseen issues, eight additional participants were recruited, resulting in a final sample of 38 participants. The selected effect size was informed by our previous work examining EVS-center-of-pressure coherence and gain (19). Although that study observed very large effects (Cohen’s *d* ≈ 2), a more conservative effect size of *d* = 1.0 was used for the original study design.

Young adults (18–35 yr old) with no self-reported neurological or orthopaedic issues that affect standing balance control were included. All participants gave written informed consent before their participation in the experiments, and all methods were reviewed and approved by the University of Waterloo Research Ethics Board (REB #47239).

### Experimental Protocol

The data analyzed in the present study were collected as part of a larger experimental protocol taking about 2 hours that also included measures of vestibulomotor responses to electrical vestibular stimulation (EVS). These vestibulomotor results have been reported separately (14). The current analyses focus on perceptual sensory weighting derived from the OCHART, as well as their relationship to vestibulomotor measures obtained from the same participants during the same experimental session.

Briefly, vestibulomotor responses were quantified using stochastic electrical vestibular stimulation (EVS) delivered during standing (14). EVS (**±** 4.5 mA) was applied as a binaural bipolar stochastic stimulus, and ML-CoP responses were recorded to assess vestibulomotor coupling. Coherence was calculated to quantify the proportion of ML-CoP variance explained by EVS, representing vestibulomotor coupling. Root mean square (RMS) of ML-CoP was also calculated to capture sway variation.

Quiet-standing EVS trials were conducted both before and immediately after the VR exposure. Each trial lasted 120 seconds, with continuous EVS throughout. Participants were instructed to stand quietly on the force plate (model 4060-05; Bertec, Columbus, OH) with their feet shoulder-width apart, arms relaxed at their sides, and eyes fixated on a taped ‘X,’ subtending a visual angle of ∼0.76°.

Participants also completed the OCHART task before and after VR, following the quiet standing EVS trials. The task was performed in two body orientations: seated upright and right side–lying, with the head supported to maintain orthogonality to gravity (Fig. 1A and B). Participants viewed the stimuli on a computer screen through a black circular shrouding tube that obscured all peripheral vision. During each 500-ms presentation, they identified whether the character was a “p” or “d” by pressing the corresponding button on a gamepad, completing a total of 1,344 trials (Fig. 1C). The character rotated around its center (coincident with a fixation point) across 24 static orientations from 0–345° in 15° increments. When displayed as a “p,” the symbol subtended 3.1° × 1.9° of visual arc; the fixation spot subtended 0.45°. Each character was displayed on either a gray background or a polarized background image, originally developed by Dyde et al. (2006), presented at one of six orientations (0–300° in 60° increments). The OCHART required approximately 15 minutes per body orientation (30 minutes total).

**Figure 1.**
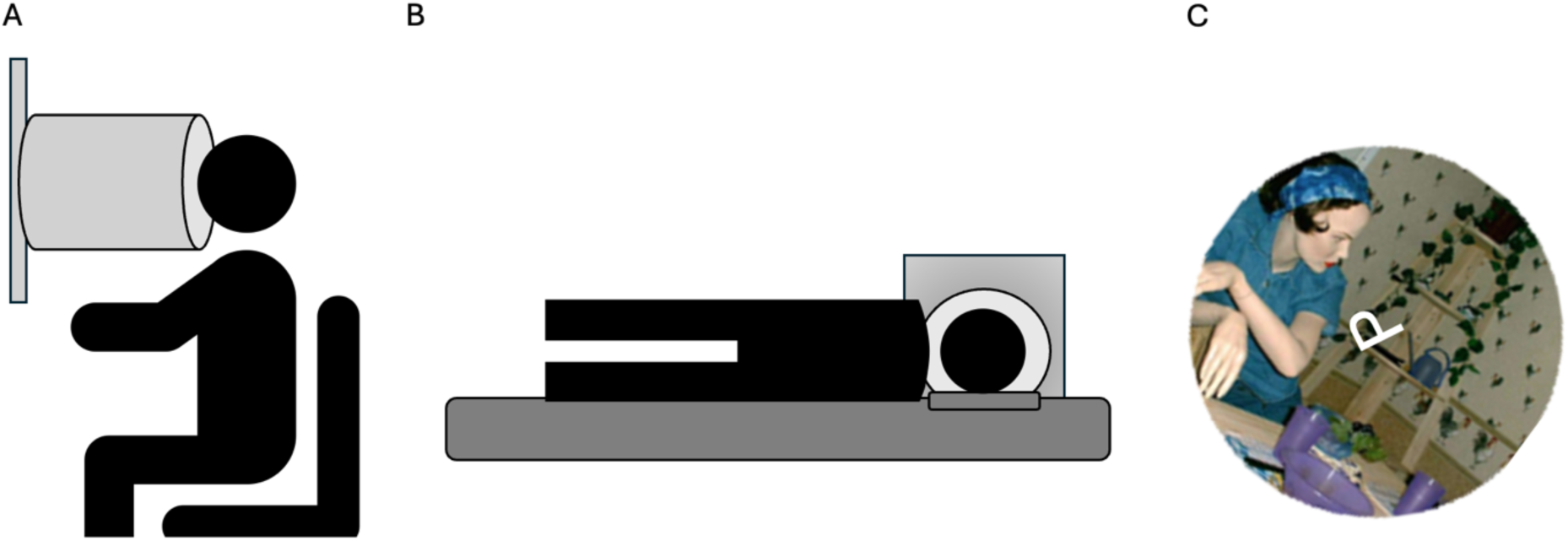
Experimental setup for the OCHART task administered before and after VR exposure. (A) Seated upright condition in which participants viewed stimuli through a black circular shrouding tube that obscured peripheral vision while maintaining fixation on a central point. (B) Right side–lying condition with the head supported to preserve orthogonality to gravity. (C) Example of a trial presented on the computer screen, showing the character at an ambiguous orientation on the polarized background image (15) at a given orientation.

After completing the quiet-standing EVS trials, participants donned the Meta Quest 1 VR system, which provides dual OLED displays (1,600 × 1,440 pixels per eye; 72 Hz refresh). Participants stood on the force plate with continuous EVS applied throughout the VR exposure, following the same postural instructions as during the quiet-standing EVS trials, except that they held the VR controllers at their sides. Participants were exposed to a highly nauseogenic rollercoaster simulation (*Epic Roller* C*oasters*; Meta Inc.) for up to 20 minutes or until they voluntarily ended the session due to discomfort. Participants completed 8 consecutive repetitions of the same rollercoaster ride, with each ride lasting approximately 2.5 minutes and no breaks between successive rides. Each repetition was identical, with the same speed, visual motion profile, and sequence of events for all participants. The visual scene incorporated translational and rotational visual motion along and about the x-, y-, and z-axes, producing a complex optic-flow stimulus throughout the ride. The VR content was passive, meaning no user input or head movement was required to progress through the simulation. A spotter provided safety support by lightly steadying a participant at the shoulder if sway approached a potential loss of balance.

Cybersickness was rated every 2.5 minutes (i.e., end of each rollercoaster ride) using the Fast Motion Sickness (FMS) Scale (2), a verbal 20-point scale ranging from 0 (“no sickness”) to 20 (“severe sickness”). Prior to starting the VR game, participants were informed that an FMS score above 15 served as a checkpoint at which discontinuing the VR exposure would be considered, although the final decision remained theirs. They were encouraged to continue if they felt comfortable doing so but were explicitly informed that choosing to stop the VR exposure would not affect their compensation and that the data collected up to that point would remain valuable for the study. Participants who discontinued the VR exposure were still able to complete the post-VR quiet-standing EVS trial and the OCHART task. These instructions were intended to minimize any perceived pressure to continue and reduce potential bias in self-reported nausea ratings.

### Sickness Groups

Participants were categorized into non-sick (FMS < 5), medium-sick (FMS ≥ 5 who completed the VR exposure), and high-sick (participants who voluntarily terminated the VR exposure early). These group definitions were established *a priori* before data collection. An FMS score < 5 was selected to identify participants with minimal symptoms, whereas participants with FMS scores ≥ 5 who completed the VR exposure were classified as medium-sick. In contrast, the high-sick group was defined by voluntary termination of the VR exposure rather than by an FMS threshold, as intolerance-driven cessation represents a distinct functional outcome beyond symptom severity alone. Defining the high-sick group using a peak FMS threshold (e.g., >15) would have excluded participants who terminated the VR exposure before reaching that rating despite being unable or unwilling to continue. Conversely, some participants exceeded an FMS score of 15 but were still able to complete the VR exposure. Therefore, voluntary discontinuation was considered a more meaningful indicator of severe cybersickness than peak symptom rating alone. This general approach is consistent with previous cybersickness research that has categorized participants into clinically meaningful symptom severity groups using symptom questionnaires, although the present thresholds were defined specifically for the FMS and the objectives of the current study (20).

This grouping approach was also appropriate for the coherence analyses conducted on the vestibulomotor responses as a part of the larger experimental protocol, which generated group-level frequency-domain estimates by concatenating the time-series data across participants within each sickness group for each condition (14). Additionally, regression approaches using FMS as a continuous variable assume linear relationships and equal intervals between ratings; however, the FMS is an ordinal scale, where differences between scores (e.g., 2–4 vs. 10–12) do not necessarily represent equivalent perceptual changes in sickness.

### OCHART Statistical Analyses

Perceptual upright (PU), defined as the orientation at which participants perceived the character as “right side up,” was estimated for each body orientation before and after VR exposure (15–18). OCHART statistical analyses were conducted using SigmaPlot 12.5. Psychometric functions were fitted to the proportion of “p” responses as a function of character orientation using a sigmoid function (eq. 1).

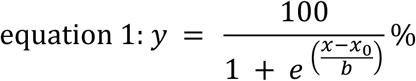

In this equation, *y* represents the percentage of “p” responses, *x₀* corresponds to the 50% point (i.e., the transition between “p” and “d” responses) and is referred to as the Point of Subjective Equality (PSE), and *b* reflects the slope of the function. Discrimination precision was quantified using the slope parameter (*b*), which reflects the width of the transition region in degrees. Smaller *b* values indicate steeper transitions and therefore greater precision. Consistent with previous OCHART studies, *b* is reported as a Just Noticeable Difference (JND), representing the smallest detectable change in stimulus orientation.

For each condition, two PSEs were identified, corresponding to the p-to-d and d-to-p response boundaries. The perceptual upright was calculated as the midpoint between these two transition angles, resulting in one PU estimate for each visual background within each body orientation (eq. 2). This led to 7 perceptual uprights for each body orientation corresponding with 6 visual backgrounds and 1 grey background.

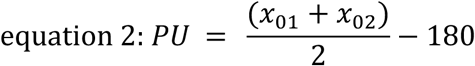

To quantify the relative contributions of visual, body, and gravity cues to perceived upright, a vector sum model was applied. In this framework, each sensory cue is represented as a unit vector oriented in the direction specified by that cue, with its contribution to the perceived upright determined by a corresponding weighting coefficient (Eq. 3). The perceived upright corresponds to the direction of the resultant weighted vector.

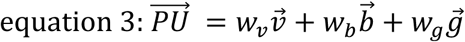

In this model, *v⃗*, *b⃗*, and *g⃗* are unit vectors representing the visual, body, and gravity cues, respectively, and *w_v_*, *w_b_*, and *w*_g_are their corresponding weighting coefficients. The gravity vector was fixed at 0° with a weighting coefficient of 1, while the visual and body weighting coefficients were optimized using a Levenberg–Marquardt algorithm to best fit the observed perceived upright values across visual backgrounds and body orientations (21).

Relative cue weights were calculated by dividing each weighting coefficient by the sum of all weighting coefficients, yielding the percentage contributions of the visual, body, and gravity cues. Changes in cue weighting were assessed by computing the difference between post- and pre-VR percentage weights for each modality. To quantify overall reweighting independent of direction, the sum of absolute changes in visual and body weights was calculated. Vector model fits were performed at the individual level, and group-level estimates were derived from averaged perceptual upright values across participants within each sickness group.

### Group Level Statistical Analyses

All analyses were performed in MATLAB (R2021b). Normality was assessed using the Shapiro–Wilk test, with p < 0.05 indicating a significant deviation from normality. Effect sizes are reported as partial eta-squared (ηp²) for ANOVAs. For non-parametric Kruskal–Wallis tests, effect sizes were calculated using eta-squared (η²).

A linear mixed-effects model was used to examine the effects of visual background orientation, time (pre- vs. post-VR), and sickness group on perceptual upright. Visual background orientation and time were included as fixed within-subject factors, sickness group was included as a fixed between-subject factor, and participant was included as a random intercept to account for repeated measurements. Significant main effects and interactions were followed by post hoc analyses where appropriate.

To determine whether there were statistical differences between sensory cue weightings of the sickness groups, a Kruskal-Wallis was conducted on the cue weights pre-and post-VR between the groups. A similar analysis was conducted to determine whether there were statistical differences between the changes in sensory cue weightings from pre-to post-VR between the sickness groups. To examine whether the magnitude of sensory reweighting was associated with sickness severity, a linear regression was performed between absolute vector-length change and peak FMS scores. Absolute change was used to capture the total magnitude of reweighting irrespective of direction. A regression approach was selected since absolute sensory reweighting represents a continuous measure that may vary across the full spectrum of sickness severity

To determine whether vestibulomotor changes were statistically related to changes in perceptual gravity cue weights, a linear regression was conducted between changes in mean coherence from pre-to-post quiet standing EVS trials and changes in gravity cue weights pre-to post-VR for each participant. This was also done for mean coherence in the pre-VR quiet standing trial and pre-VR gravity weights. To determine whether sway responses experienced during VR were statistically related to perceptual visual cue weights, a simple regression was conducted between total ML-CoP RMS during the VR trial and pre-VR visual cue weights.

## Results

### Post Collection Inclusion/Exclusion

Three participants (one from each sickness group) were excluded because the vector sum model could not be reliably fit to their data. In these cases, participants did not demonstrate clear points of subjective equality, with most character orientations showing ambiguous responses over many trials (i.e., neither clearly “P” nor “d”). As a result, perceptual upright values could not be estimated. This ambiguity may reflect variability in task performance or difficulty resolving perceptual upright.

Spotter assistance was required for three participants, all in the high-sick group. In each case, the spotter lightly stabilized the participant’s shoulders to prevent loss of balance; however, these data were retained for analysis.

### Sickness

Using the *a priori* classification criteria described in the methods, participants were categorized into non-sick, medium-sick, and high-sick groups based on their cybersickness responses during the VR rollercoaster task. The non-sick group (peak FMS < 5) had 12 participants, the medium-sick group (peak FMS ≥ 5 who completed the VR exposure) had 13 participants, and the high-sick group (participants who voluntarily discontinued the VR exposure before 20 minutes) had 13 participants. The group-specific FMS trajectories during the VR exposure were identical to those reported in our previous study using the same participant cohort and are therefore not reproduced here (14).

### OCHART

To determine whether visual background orientation influenced PU judgments differently after VR exposure and across sickness groups, we examined how PU changed across visual background orientations pre-and post-VR and across sickness groups. A linear mixed-effects analysis revealed a significant main effect of visual background orientation (*F* (5, 384) = 11.93, *p* < 0.001), indicating that perceptual upright varied across visual background orientations (Fig. 2). No significant main effects of time (*F* (1, 384) = 0.21, *p* = 0.648) or sickness group (*F* (2, 384) = 0.24, *p* = 0.784) were observed. Furthermore, there were no significant interactions between time and visual background orientation (*F* (5, 384) = 0.06, *p* = 0.997), time and sickness group (*F* (2, 384) = 0.01, *p* = 0.991), visual background orientation and sickness group (*F*(10, 384) = 1.23, *p* = 0.268), or the three-way interaction (*F* (10, 384) = 0.61, *p* = 0.804). These findings indicate that the influence of visual background orientation on perceptual upright remained stable following VR exposure and did not differ across sickness groups.

**Figure 2.**
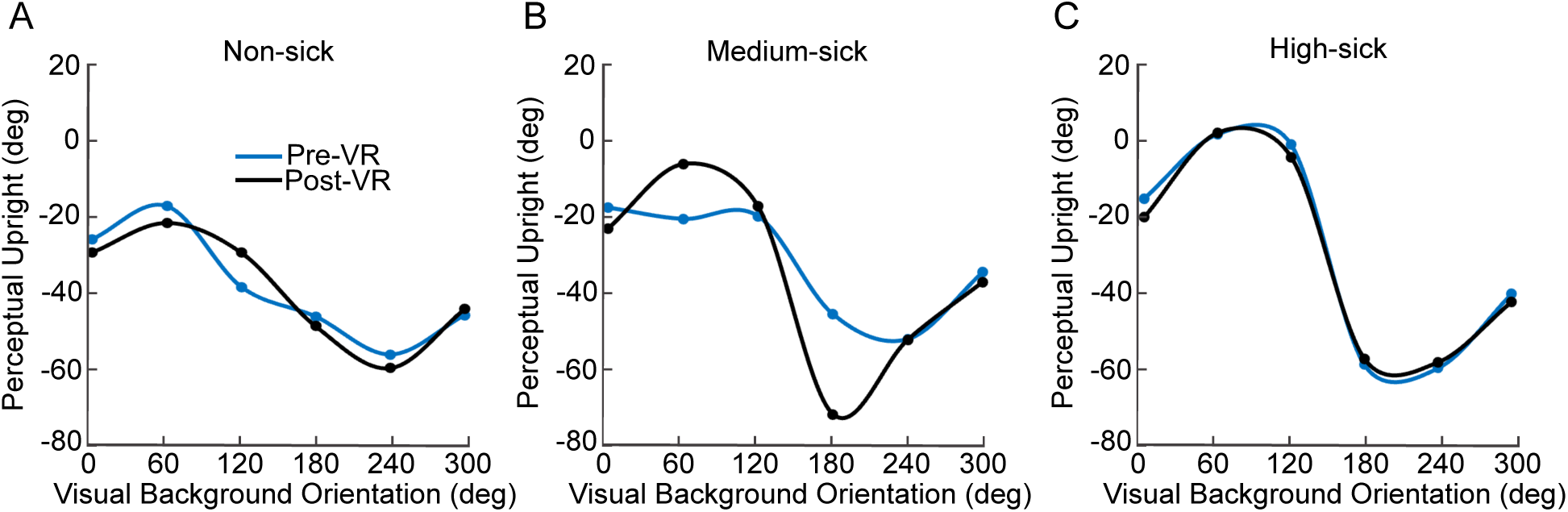
Mean perceptual upright values across visual background orientations during right-side-down trials before (blue) and after (black) VR exposure for the non-sick (left), medium-sick (middle), and high-sick (right) groups.

Estimates of group-level cue weights were derived from averaged perceptual upright values across participants within each sickness group. These estimates indicated that non-sick individuals exhibited greater reliance on gravity cues and lower reliance on visual cues, whereas highly sick individuals showed lower gravity weights and higher visual weights. Specifically, prior to VR exposure, the non-sick group weighted body, gravity, and vision cues at 46%, 36%, and 18%, respectively, and showed similar weighting post-VR (46% body, 35% gravity, 19% vision; Fig. 3). The medium-sick group exhibited cue weights of 50% body, 31% gravity, and 19% vision pre-VR, which shifted to 43% body, 31% gravity, and 26% vision post-VR. In the high-sick group, cue weights were 45% body, 25% gravity, and 30% vision pre-VR and remained comparable post-VR (44% body, 26% gravity, 30% vision).

**Figure 3.**
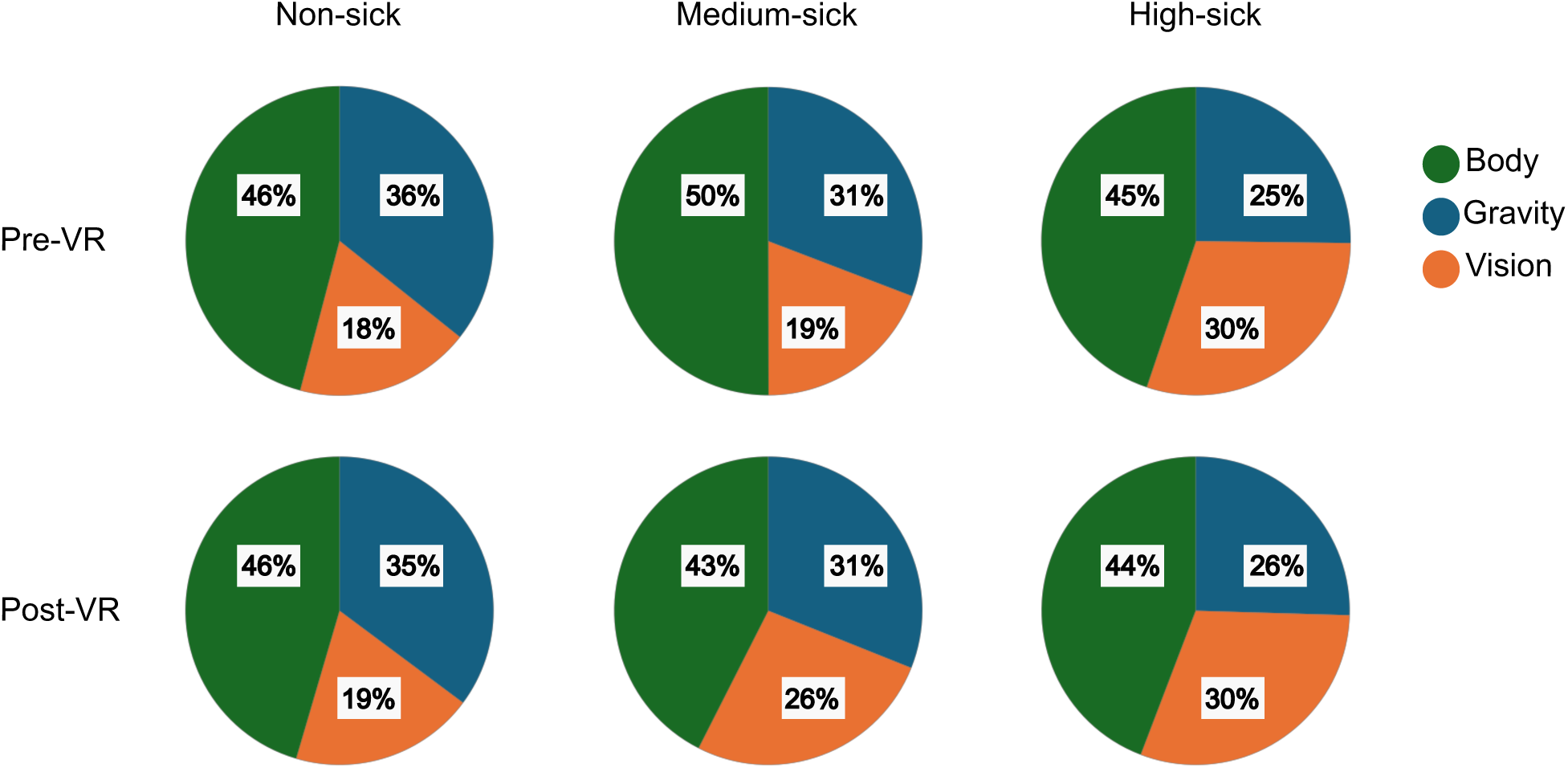
Group-level perceptual cue weights derived from the OCHART perceptual upright task before and after VR exposure. Pie charts show the relative weighting of body (green), vision (orange), and gravity (blue) cues for the non-sick (left), medium-sick (middle), and high-sick (right) groups. Pre-VR weights are shown in the top row, and post-VR weights are shown in the bottom row. Percentages represent mean cue weights averaged across participants within each group, with sample sizes indicated above each column.

Statistical differences in individual perceptual cue weights between sickness groups were also examined. A Kruskal–Wallis test of individual perceptual cue weights revealed a significant group difference in gravity weights before VR exposure (χ² (2) = 6.29, *p* = 0.043, η² = 0.134; Fig. 4A). Post hoc Tukey tests indicated that non-sick participants had higher gravity weights than high-sick participants (*p* = 0.034). No other between-group differences were observed. Specifically, gravity weights did not differ significantly after VR exposure (ANOVA: *F* (2, 32) = 2.56, *p* = 0.101; Fig. 4D), nor did vision weights before (Kruskal–Wallis: χ² (2) = 2.72, *p* = 0.181; Fig. 4B) or after VR exposure (ANOVA: *F* (2, 32) = 1.10, *p* = 0.345; Fig. 4E), or body weights before (ANOVA: *F* (2, 32) = 0.504, *p* = 0.609; Fig. 4C) or after VR exposure (ANOVA: *F* (2, 32) = 0.08, *p* = 0.926; Fig. 4F).

**Figure 4.**
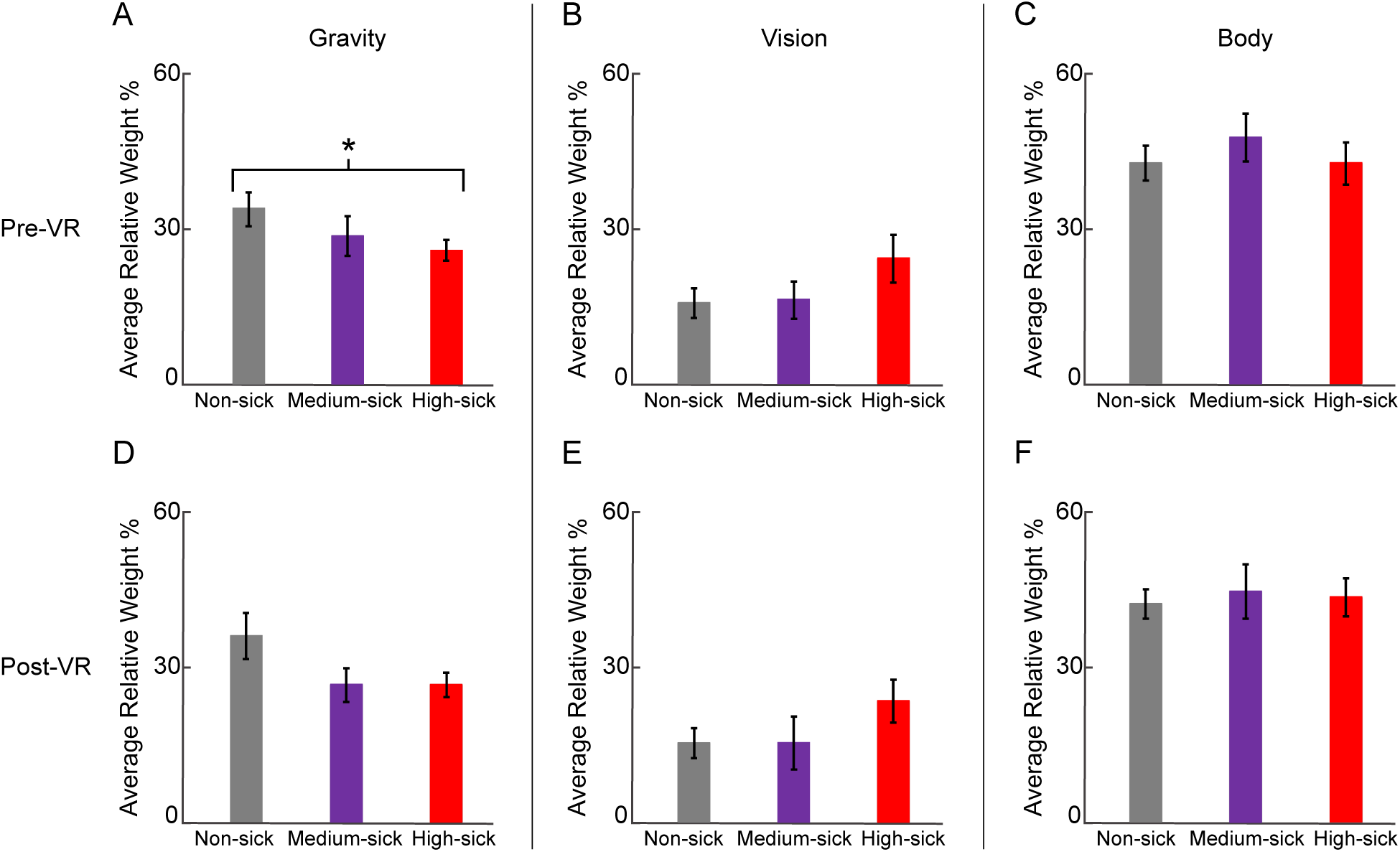
Individual perceptual cue weights derived from the OCHART perceptual upright task across sickness groups before and after VR exposure. Bar plots show relative weighting of gravity (A, D), vision (B, E), and body (C, F) cues for the non-sick, medium-sick, and high-sick groups. Pre-VR cue weights are shown in the top row (A–C), and post-VR cue weights are shown in the bottom row (D–F). Bars represent group means and error bars indicate ± SEM. Asterisks denote statistically significant between-group differences (*p* < 0.05).

Although additional between-group differences beyond pre-VR gravity weights were not statistically significant, the direction of effects was consistent with overall group-level trends. Before VR exposure, gravity weights were highest in the non-sick group (38 ± 4%), intermediate in the medium-sick group (32 ± 4%), and lowest in the high-sick group (29 ± 2%; Fig. 4A). A similar pattern was observed following VR exposure, with gravity weights remaining highest in the non-sick group (38 ± 5%) and lower in both the medium-sick (28 ± 4%) and high-sick groups (28 ± 3%; Fig. 4D). Vision weights showed the opposite pattern. Before VR exposure, visual weighting was lowest in the non-sick group (16 ± 3%), slightly higher in the medium-sick group (17 ± 4%), and highest in the high-sick group (26 ± 5%; Fig. 4B). Similar values were observed following VR exposure, with vision weights of 16 ± 3% in the non-sick group, 16 ± 5% in the medium-sick group, and 25 ± 4% in the high-sick group (Fig. 4E). Body weights varied minimally across groups and conditions. Before VR exposure, body weights were 45 ± 4% in the non-sick group, 51 ± 5% in the medium-sick group, and 45 ± 4% in the high-sick group (Fig. 4C). Following VR exposure, body weights remained similar across groups at 45 ± 3%, 48 ± 6%, and 46 ± 4%, respectively (Fig. 4F).

Statistical differences in changes in individual perceptual cue weights between sickness groups were examined. Changes in individual perceptual cue weights from pre- to post-VR exposure were small, variable, and did not differ significantly across sickness groups. For gravity weights, changes were 0.44 ± 2.48% in the non-sick group, −3.74 ± 3.25% in the medium-sick group, and −1.69 ± 1.98% in the high-sick group (Kruskal–Wallis: χ² (2) = 0.54, *p* = 0.764; Fig. 5A). For vision weights, changes were −0.12 ± 1.03%, 6.88 ± 5.02%, and −0.64 ± 1.54% in the non-sick, medium-sick, and high-sick groups, respectively (Kruskal–Wallis: χ² (2) = 0.08, *p* = 0.959; Fig. 5B). For body weights, changes were −0.32 ± 2.75%, −3.14 ± 2.47%, and 2.33 ± 1.67% in the non-sick, medium-sick, and high-sick groups, respectively (Kruskal–Wallis: χ² (2) = 3.89, *p* = 0.243; Fig. 5C).

**Figure 5.**
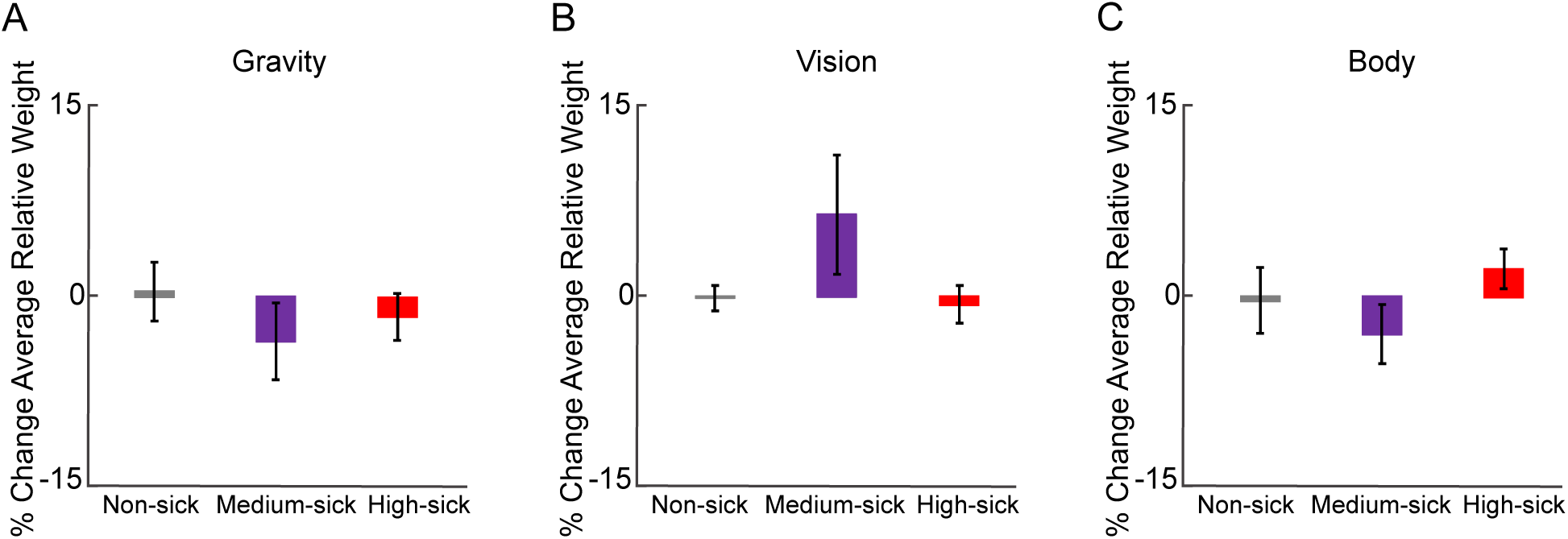
Percent change in individual perceptual cue weights from pre- to post-VR exposure across sickness groups. Bar plots show percent change in gravity (A), vision (B), and body (C) cue weights for the non-sick, medium-sick, and high-sick groups. Bars represent group means and error bars indicate ± SEM.

To determine whether overall perceptual reweighting scaled with symptom severity rather than categorical sickness group, we examined the relationship between absolute sensory reweighting magnitude and peak FMS score. A linear regression relating absolute vector-length change of the perceptual upright (the sum of visual and body cue changes) to peak FMS score revealed no significant association, indicating that total sensory reweighting magnitude, irrespective of direction, was not related to sickness severity (*p* = 0.746, R² = 0.003; Fig. 6A). Even when outlier participants were removed (change in vector length > 2), there was no significant association (*p* = 0.112, R² = 0.085; Fig. 6B).

**Figure 6.**
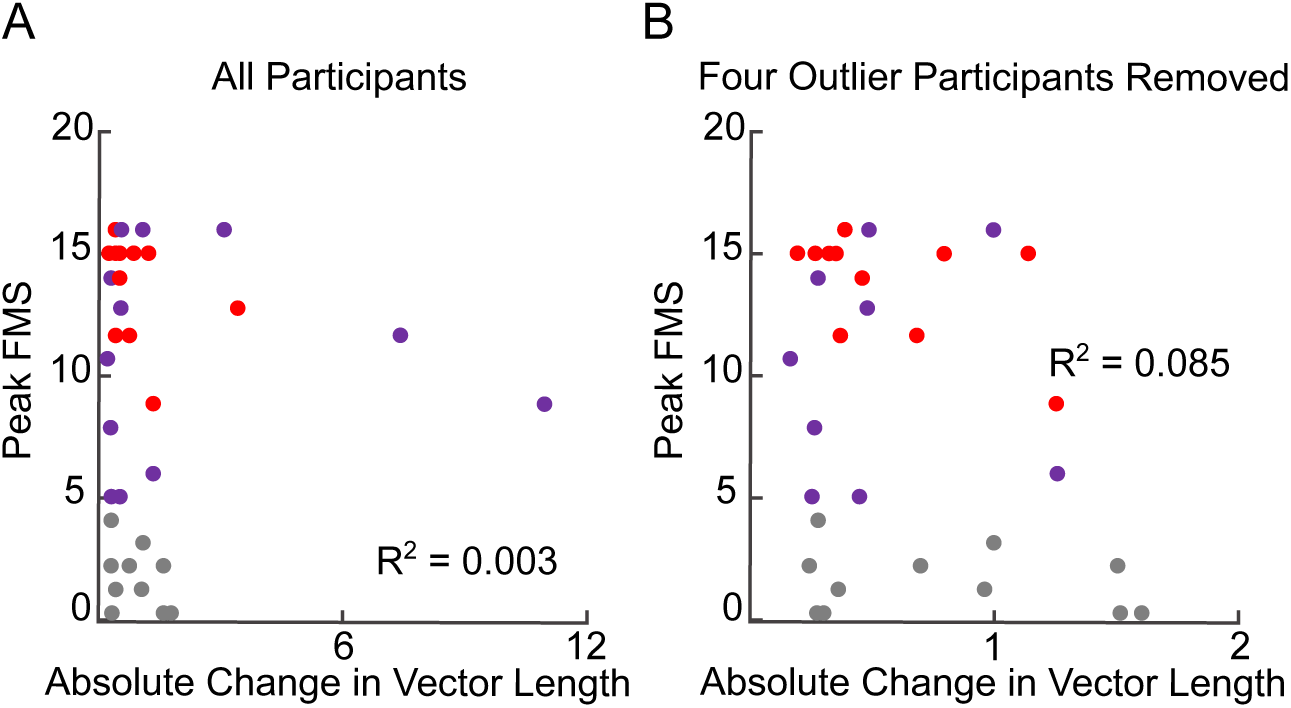
Relationship between peak sickness severity and absolute sensory reweighting magnitude. (A) Linear regression between peak Fast Motion Sickness (FMS) score (y-axis) and absolute vector-length change (x-axis), calculated as the sum of visual and body cue weight changes from pre- to post-VR. (B) The same relationship shown after removal of four outlier participants, defined as those with an absolute vector-length change exceeding 2. Each point represents an individual participant and is coloured according to sickness group (grey, non-sick; purple, medium-sick; red, high-sick). R² values are shown for each regression.

A linear regression relating changes in mean EVS-ML-CoP coherence from pre- to post-VR quiet-standing EVS trials (vestibulomotor changes) to changes in gravity cue weights from pre- to post-VR exposure (perceptual changes) revealed no significant association across participants (*p* = 0.410, R² = 0.021; Fig. 7A). Similarly, there was no significant association between pre-VR vestibulomotor coherence and pre-VR relative gravity weight (*p* = 0.592, R² = 0.009; Fig. 7B), or between total ML-CoP RMS during the VR trial and pre-VR visual cue weights (*p* = 0.279, R² = 0.035; Fig. 7C)

**Figure 7.**
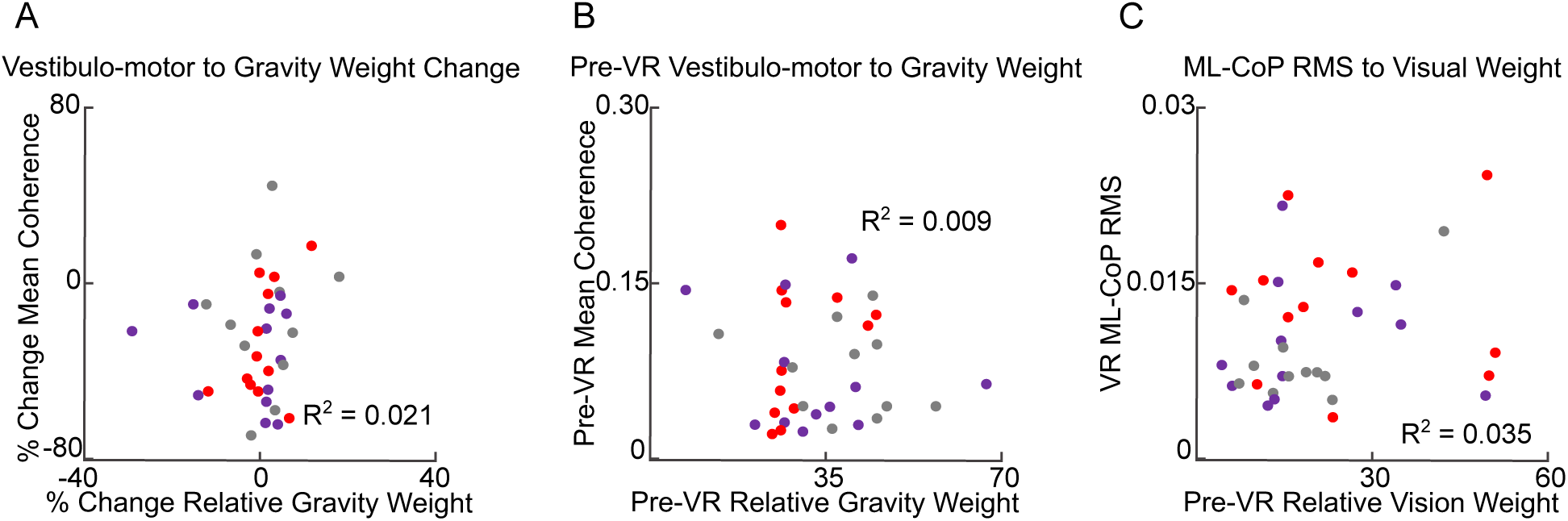
Relationships between sensory weighting and vestibulomotor or postural responses. (A) Association between percent change in vestibulomotor coherence and percent change in relative gravity weight from pre- to post-VR exposure. (B) Relationship between pre-VR relative gravity weight and pre-VR vestibulomotor coherence. (C) Association between pre-VR relative visual weight and mediolateral center-of-pressure root mean square (ML-CoP RMS) during VR exposure. Each point represents an individual participant and is coloured according to sickness group (grey, non-sick; purple, medium-sick; red, high-sick). R² values are shown for each regression.

## Discussion

This study examined how perceptual sensory weighting during immersive VR exposure with conflicting EVS contributes to individual susceptibility to cybersickness and whether perceptual sensory weights are related to previously reported vestibulomotor measures. Consistent with our hypotheses, greater pre-exposure reliance on gravity cues was associated with lower sickness susceptibility. Although reduced visual weighting showed a directional trend, differences between sickness groups were not statistically significant. Meaningful changes in perceptual weighting or in the influence of visual background orientation from pre- to post-VR were minimal, suggesting that stable baseline sensory weighting strategies may play a more prominent role in cybersickness susceptibility than short-term perceptual reweighting during the present VR exposure. Vestibulomotor responses were not correlated with gravity weights in perceptual orientation, and greater postural sway during VR was not associated with increased visual weights. These findings indicate that perceptual object orientation and motor postural control likely reflect partially distinct processes of sensory integration. Improving our understanding of these mechanisms may enhance our ability to predict susceptibility to cybersickness and inform targeted interventions and design strategies to promote safer and more accessible VR applications across education, healthcare, and industry.

### Non-Sick Individuals Exhibit Greater Gravity Weighting During Orientation Perception

Our findings show that non-sick individuals exhibited greater gravity cue weights both before (36% vs. 25%) and after VR exposure (35% vs. 25%) compared with the high-sickness group. Conversely, the high-sickness group demonstrated higher visual weights than the non-sick group (30% vs. 18% pre-VR and 30% vs. 19% post-VR); however, these differences did not reach statistical significance. Despite this directional trend, only pre-VR gravity weighting differed significantly between groups. This finding suggests that gravity weighting may represent a more sensitive perceptual marker of cybersickness susceptibility than visual weighting in the present paradigm. The observed effect was modest and evident only between the non-sick and high-sick groups. Even so, these findings are consistent with our hypothesis that greater reliance on gravity cues is associated with reduced susceptibility to cybersickness and with previous work suggesting that greater gravity weighting supports stable orientation perception under conditions of visual ambiguity (11).

Previous studies using the OCHART paradigm have shown that perceived upright is typically dominated by the egocentric body cue, with the body weighting accounting for over half of the total weighting, while vision and gravity contribute smaller proportions (54% body, 25% vision and 21% gravity for the PU (15,17)). In the present study, the non-sick group exhibited higher estimated gravity weighting before VR exposure (46% body, 18% vision, and 36% gravity), whereas the high-sick group showed a shift toward greater visual weighting (45% body, 30% vision, and 25% gravity). Notably, gravity weighting was higher in both groups than has been reported previously. One possible explanation is that the addition of EVS altered the relative weighting of sensory cues during the perceptual upright task. Since EVS introduces artificial vestibular signals that differ from natural vestibular input, the resulting gravity weights may not be directly correspond to values reported in studies performed without vestibular perturbation.

Greater reliance on gravity cues may provide a more stable reference frame against which conflicting visual motion can be evaluated. During the rollercoaster simulation, participants were exposed to compelling visual signals of self-motion that were not matched by corresponding vestibular and somatosensory inputs. Individuals who place greater weight on gravity-related cues may be less likely to accept visually specified motion as veridical, instead maintaining orientation estimates anchored to an internal representation of upright relative to gravity. In contrast, individuals who rely more heavily on visual information may be more susceptible to adopting the visually specified motion, increasing the discrepancy between visual and non-visual sensory signals. This larger sensory conflict could contribute to the development of cybersickness symptoms. From this perspective, greater gravity weighting may not directly reduce sickness but instead reflect one aspect of a sensory integration strategy that is more resistant to visually induced sensory conflict.

### Perceptual Cue Weighting Remained Stable Following VR Exposure

Perceptual cue weights showed little change from pre- to post-VR across all sickness groups, in contrast to previous findings of perceptual reweighting (11). Consistent with this finding, the influence of visual background orientation on perceptual upright remained stable following VR exposure and did not differ across sickness groups, suggesting that contextual visual influences on orientation perception were likewise unaffected by the VR exposure. This relative stability suggests that, in the present paradigm, baseline perceptual weighting may play a more prominent role than short-term adaptation in determining susceptibility to cybersickness. One possible explanation is that passive VR exposure imposes greater demands on postural control than on perceptual orientation judgments, limiting the extent to which perceptual reweighting occurs over the time course examined.

### Dissociation Between Sensorimotor and Perceptual Outcomes

Vestibulomotor responses were not correlated with gravity-based weights derived from perceptual orientation judgments. Likewise, postural sway magnitude during VR exposure was not related to participants’ reliance on visual cues when determining perceptual upright. Together, these findings suggest that perceptual orientation judgments and postural control may rely on partially independent multisensory integration processes. Specifically, the use of gravitational references for orientation perception may be dissociable from the use of vestibular inputs for balance control. Similarly, visual cue use during perceptual orientation judgments may not directly correspond to visually induced balance responses. One possible explanation is that visual weighting during perceptual orientation reflects how visual information contributes to conscious estimates of upright, whereas ML-CoP RMS reflects the integrated motor output of multiple sensory and biomechanical processes during dynamic balance control. Consequently, greater reliance on visual cues for orientation perception may not necessarily translate to larger visually induced sway responses during VR exposure.

This dissociation may also reflect differences in task sensitivity between sensorimotor and perceptual domains. The experimental paradigm placed substantial demands on balance control by perturbing both vestibular and visual inputs, whereas perceptual judgments may have been comparatively less challenged. This imbalance in task demands may help explain why reweighting was observed at the vestibulomotor level but not in perceptual cue weighting, and why perceptual visual weighting did not predict visually induced postural sway.

The lack of correspondence between vestibulomotor responses and perceptual gravity weighting may also reflect fundamental differences in how these measures are derived and the neural processes they capture. Vestibulomotor responses provide a relatively direct index of how vestibular information contributes to postural control by quantifying the relationship between experimentally imposed vestibular perturbations and motor output. In contrast, OCHART-derived gravity weights are inferred through a static vector-sum model of perceptual upright. The resulting estimates do not isolate vestibular processing per se but instead reflect the net outcome of multiple interacting sensory and central processes that determine how visual, body, and gravitational references are integrated to form a conscious orientation judgment.

Consequently, a higher “gravity weight” in the OCHART should not necessarily be interpreted as greater vestibular sensitivity or increased reliance on vestibular afferent input. Rather, OCHART-derived gravity weights may reflect stable individual differences in how the CNS constructs and maintains an internal representation of upright orientation, including the extent to which individuals rely on gravitational information when visual and body cues conflict. Such processes may involve higher-order cortical networks implicated in spatial orientation and multisensory integration, in addition to the brainstem pathways supporting rapid vestibulomotor responses.

The observed dissociation between perceptual and vestibulomotor measures may therefore reflect a meaningful distinction rather than simply a methodological difference. Individuals who maintain stable perceptual representations of upright despite conflicting visual input may be protected from cybersickness through mechanisms distinct from those governing online balance control. Thus, perceptual estimates of upright and vestibulomotor responses may provide complementary information about how the CNS resolves sensory conflict, operating over different timescales and through partially distinct neural substrates.

### Limitations and Future Directions

Several limitations should be considered when interpreting the present findings. First, the study employed EVS with a peak amplitude of ±4.5 mA to probe vestibulomotor contributions to balance during VR exposure. While EVS provides a controlled method for quantifying vestibular influence, it introduces high-intensity and non-physiological vestibular signals that likely increase cybersickness. As such, the OCHART responses are influenced by perturbed, rather than natural, vestibular input.

Second, participant grouping was based on categorical classifications of sickness severity and task intolerance rather than continuous measures. While this approach captures meaningful functional differences (particularly the distinction between individuals who completed the task and those who terminated early) and facilitates frequency-dependent concatenation analyses required in the original experimental protocol, it may obscure more subtle relationships between symptom progression and sensory reweighting.

Third, perceptual sensory weighting was assessed using a static perceptual upright task administered before and after VR exposure. Although OCHART is a well-validated measure of orientation perception, it may not be sufficiently sensitive to detect short-term perceptual adaptations during a highly dynamic VR experience. OCHART assumes that perceptual upright can be decomposed into relatively stable contributions from gravity, visual, and body cues. Consequently, OCHART-derived weights should be interpreted as estimates of relatively stable perceptual biases rather than direct measures of moment-to-moment sensory reweighting. More immersive or continuous perceptual measures, such as real-time judgments of verticality during VR exposure, may better capture transient perceptual adaptations and clarify how sensory estimates are updated during ongoing sensory conflict. Combining such measures with physiological indices of vestibular processing may further bridge the gap between perceptual and vestibulomotor domains.

Fourth, the VR paradigm placed substantial demands on postural control, particularly during visually salient perturbations such as the initial rollercoaster drop. In contrast, perceptual orientation judgments were performed in a comparatively less challenging context. This imbalance in task demands may have contributed to the observed dissociation between sensorimotor and perceptual outcomes. Future work could systematically manipulate the difficulty of perceptual and postural tasks to better examine how sensory weighting adapts across domains under matched levels of challenge.

Finally, although the cohort was adequately powered for the primary vestibulomotor outcomes of the parent study, the present secondary analysis may have been underpowered to detect more subtle differences in perceptual weighting, particularly for visual cue weighting.

## Overall Summary

In summary, individual differences in baseline perceptual sensory weighting are associated with susceptibility to cybersickness during immersive VR exposure. Individuals who relied more heavily on gravity cues appeared better equipped to tolerate VR-induced sensory conflict. Reduced visual weighting exhibited a similar directional trend but was not significantly associated with sickness group. In contrast, perceptual reweighting showed minimal change. Importantly, perceptual and vestibulomotor measures were largely dissociable, suggesting that orientation perception and postural control rely on partially independent integration processes or differ in task sensitivity. These findings highlight the critical role of baseline sensory integration in shaping responses to sensory conflict and have important implications for predicting and mitigating cybersickness in virtual environments.

## Data Availability

All data and code for this article are provided in an Open Science Framework (OSF) repository, at https://osf.io/9n6hf. We report all measured variables and all analyses we conducted.

## Disclosures and Disclaimers

The authors have no disclosures or disclaimers to declare.

## Author Contributions

MHG developed the main concept and theoretical framework of the project. MHG designed and conducted the experiment, analyzed the data, and led the writing of the manuscript. MBC secured funding and provided oversight of the project’s direction and planning. Both authors contributed critical feedback and helped shape the research, analysis, and manuscript.

